# Altered synaptic connectivity in an *in vitro* human model of STXBP1 encephalopathy

**DOI:** 10.1101/2021.09.22.461333

**Authors:** Faye McLeod, Anna Dimtsi, David Lewis-Smith, Rhys Thomas, Gavin J Clowry, Andrew J Trevelyan

## Abstract

Early infantile developmental and epileptic encephalopathies are devastating conditions, generally of genetic origin, but the pathological mechanisms often remain obscure. A major obstacle in this field of research is the difficulty of studying cortical brain development in humans, *in utero*. To address this, we established an *in vitro* assay to study the impact of gene variants on the developing human brain, using living organotypic cultures of the human subplate and neighbouring cortical regions, prepared from ethically sourced, 14-17 post conception week brain tissue (www.hdbr.org). We were able to maintain cultures for several months, during which time, the gross anatomical structures of the cortical plate, subplate and marginal zone persisted, while neurons continued to develop morphologically, and form new synaptic networks. This preparation thus permits the study of genetic manipulations, and their downstream effects upon an intact developing human cortical network. We focused upon STXBP1 haploinsufficiency, which is among the most common genetic causes of developmental and epileptic encephalopathy. This was induced using shRNA interference, leading to impaired synaptic function and a drop in the number of glutamatergic synapses. We thereby provide a critical proof-of-principle for how to study the impact of any gene of interest on the development of the human cortex.

## Introduction

Approximately 1 in 2000 children in the UK develop epilepsy before three years of age from a monogenic cause^1^; of these, about a third will continue to have seizures into adulthood^2^. Knowing the genetic cause aids clinical management, yet the true pathological mechanisms remain obscure, at least in part because the aetiology often starts *in utero*, making their study extremely difficult. A critical phase of prenatal neocortical development is the transient appearance, and subsequent dissolution, of the subplate. This cortical region contains the earliest cortical synaptic network and forms the scaffold for subsequent neocortical development^3, 4^. It is therefore the site where synaptic malfunction will first disrupt the developing cortex.

Of particular interest is Syntaxin-binding protein 1 (STXBP1), an essential pre-synaptic protein also called MUNC18-1^5^, that is highly expressed in the prenatal developing brain^6^. Loss of function variants in *STXBP1*, collectively termed ‘STXBP1 encephalopathies’, result in significant intellectual disability without regression, early-life onset epilepsy in most, and a range of movement disorders including ataxia, tremor and spasticity^7–9^; but the underlying pathological mechanisms are poorly understood. Individuals with protein truncating variants seem to have similar clinical presentations when compared in a large cohort with STXBP1 encephalopathies, suggesting a shared pathological aetiology^10^. It seems likely that the effects of STXBP1 haploinsufficiency arise prenatally, supported by the fact that STXBP1 is highly expressed in the developing subplate^6^. The study of functional prenatal interactions, however, presents a major practical challenge for early human development.

STXBP1 haploinsufficiency has been replicated in transgenic mouse models and is associated with a postnatal deficit in synaptic neurotransmission^11, 12^. Rodent models of this condition though are limited in one important respect: the rodent subplate is small and very simple in organisation, compared with the human subplate^13^, calling into question the relevance of these rodent models to human development. Likewise, human cerebral organoids are similarly limited, because, to date, there have been no descriptions of any organoid structures that accurately resembles the subplate as it exists *in utero*^14^.

To overcome this critical gap in our experimental arsenal, we have developed a protocol for preparing living organotypic cultures of human neocortex, and its underlying subplate, from ethically approved samples of human fetal brain tissue, provided by the MRC-Wellcome Trust Human Developmental Biology Resource (HDBR, www.hdbr.org). Over time, neurons within the culture display evolving morphology, while the culture itself retains its gross anatomical structure. Importantly, the cultures are amenable to genetic manipulation, including knock-down of STXBP1. This is achieved using short hairpin RNA (shRNA) interference, delivered by adeno-associated viral (AAV) viral vectors, to mimic the disease-causing, haploinsufficient state, and by using viral vectors carrying a scrambled *STXBP1* sequence as a control. STXBP1 knock-down results in altered synaptic stability and impaired spontaneous pre-synaptic neurotransmitter release, and changes in the number of synaptic terminals. We have thus developed a model that is closer to early human neurodevelopment than existing alternatives, and able to facilitate the study of mechanisms suspected of causing neurodevelopmental disorders in humans.

## Materials and methods

Detailed methods are provided in the supplementary material.

### Human organotypic slice cultures

Human foetal brain tissue was acquired from the HDBR; (https://www.hdbr.org/) in Newcastle upon Tyne (Project 200428). Full consent was given in line with ethical approval from the Newcastle and North Tyneside NHS Health Authority Joint Ethics Committee (REC reference: 18/NE/0290). Cultures were prepared from 10 different foetal brains (three from 14 post-conception week (pcw), four 16 pcw and three 17 pcw). Slices were sectioned on a vibratome (280 μm), and then transferred to 6-transwell plates containing culture media, which was changed every two days. The plates were maintained at 37°C, 5% CO_2_ in ambient O_2_ and 90% humidity.

### Viral vectors

Three AAV vectors were used (VectorBuilder):

1. Scrambled control shRNA - packaged into an AAV9 vector expressing the cytomegalovirus (CMV) promoter and enhanced green fluorescent protein (eGFP) to achieve a final titre of 5×10^12^ GC/ml. The scrambled target sequence was 5’-CCTAAGGTTAAGTCGCCCTCG-3’. Infected cells were referred to as ‘control’.
2. STXBP1 shRNA - packaged into an AAV9 vector expressing the CMV promoter and eGFP to achieve a final titre of 5×10^12^ GC/ml. The STXBP1 target sequence was 5’TACTGAAGCACAAACATATAT-3’.
3. AAV9-CMV-GFP (final titre of 1×10^13^ GC/ml).

Infected cells were identified by their eGFP/GFP tag. At 0 days *in vitro* (DIV), 3-5 μl of viral vectors were added to the top of each slice. All experiments were conducted 1-4 weeks later.

### Live imaging

Live imaging was performed using an inverted Zeiss LSM800 Airyscan confocal microscope. FM4-64 dye was utilised as previously described^15^ and analysis was performed using Fiji software (NIH, RRID:SCR_002285).

### Whole cell-patch clamp recordings

Slices were transferred into the recording chamber of an upright Leica DMLFSA fluorescent microscope fitted with Micro Control Instruments micromanipulators, and continuously perfused at 34°C with oxygenated artificial cerebro-spinal fluid. AMPA receptor-mediated miniature glutamatergic post-synaptic currents were recorded at −60 mV in the presence of 100 nM TTX, 10 μM bicuculline, 50 μM AP-5. Miniature GABAergic post-synaptic currents were recorded at 0 mV in the presence of 100 nM TTX and 50 μM AP-5.

### Immunohistochemistry

Staining was performed using a protocol optimised specifically for thick slices^16^.

### Image acquisition and analysis

Images were acquired on a Zeiss LSM800 Airyscan confocal microscope with fixed settings. The x5 objective (NA = 0.16) was used to acquire image stacks (z-step = 5 μm) of general slice anatomy. The ×10 objective was used to aid identification of the subplate. This region has an easily identifiable anatomy with a sparse cell population in comparison to the neighbouring cortical plate. Subsequently, x20 objective (NA = 0.8) with x2 zoom was used to acquire image stacks (z-step = 2 μm) of the infected and non-infected subplate neurons and the x63 objective (NA = 1.4) to acquire image stacks (z-step = 0.25 μm) of synaptic components. Three to four image stacks in the upper layer of the subplate were acquired per slice in a randomised manner. A final image resolution of 1024×1024 pixels was obtained.

The number as well as the length of primary subplate neuron processes were measured using Fiji software. Synaptic counts and integrated density (area × mean intensity) measurements were conducted using Zen 3.1 software (Zeiss; RRID:SCR_013672).

### Western blot

Equal homogenate amounts were run on 10% SDS-PAGE gel and transferred onto PVDF membranes using BIO-RAD western blot apparatus.

### Statistical analysis

Statistical analyses were performed on GraphPad Prism 9 (RRID:SCR_002798). We assessed data normality using Kolmogorov-Smirnov tests. The appropriate parametric test was used for all normally distributed data, including paired/unpaired t-tests, one-way ANOVA and two-way ANOVA with Tukey’s multiple comparisons test and repeated measures. Statistical significance was accepted as * p < 0.05, ** p < 0.01, and *** p < 0.001, and non-significant data were indicated as ns. All data in the results section are presented as ± SEM. Further details of exact sample replicates and statistical tests used are stated within the results section and each figure legend.

### Data availability

All data generated or analysed and used in this study are available upon reasonable request.

## Results

### STXBP1 expression in the developing human brain

We created organotypic cortical slice cultures from fresh human foetal brain tissue samples from 10 different individuals. At the time of preparation, the *in utero* developmental age of samples was 14–17 pcw. In all cases, the cultures retained the distinct and identifiable anatomical regions of the subplate, its neighbouring cortical plate, and the marginal zone (Figure 1A-B). Synapsin1 immunoreactivity revealed that pre-synaptic terminals were predominantly located in the upper layer of the subplate, at this stage of development (Figure 1A). The subplate was smallest in 14 pcw samples (n = 3 individual samples; average depth = 308 ± 57 μm), becoming progressively larger in the cultures prepared from 16 (n = 4; 525 ± 11 μm) and 17 pcw (n = 3; 651 ± 33 μm) (p < 0.05, one-way ANOVA; Figure 1B). STXBP1 showed higher expression at 16-17 pcw, compared to 14 pcw (Figure 1B), and was observed near spine-like structures and along putative axons of subplate neurons (Figure 1D). Collectively these data show that the subplate becomes more prominent and enriched with pre-synaptic proteins including STXBP1 over the 14–17 pcw period.

**Figure 1.**
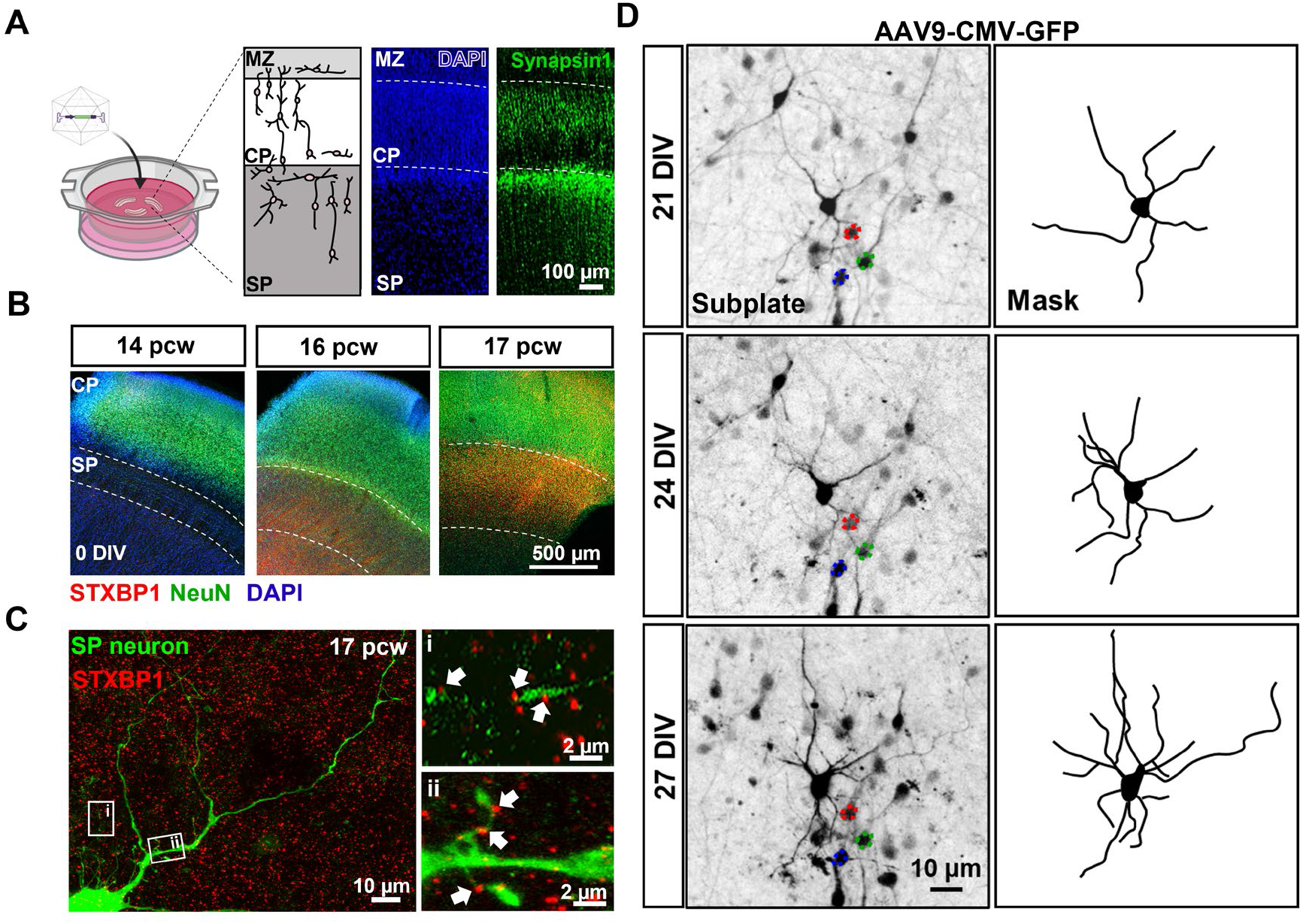
Human organotypic cortical brain slices cultures with developing subplate neurons. (**A**) Schematic and low magnification photomicrographs of organotypic cultures prepared from human foetal brain tissue. Cultures are maintained for 1–4 weeks following infection with and without different viral vectors. Micrograph image represents a 16 pcw sample, 2 weeks in culture with clearly identifiable cortical regions including the marginal zone (MZ), cortical plate (CP) and subplate (SP). DAPI labels all cell nuclei and Synapsin1 immunohistochemistry labels reveals pre-synaptic terminals predominantly located in SP but also CP and MZ. **(B)** Representative images from 14, 16 and 17 pcw cultures (0 DIV). STXBP1 (red) and NeuN (green, all neurons) immunofluorescence reveals location of the SP. Note the enrichment of STXBP1 within the SP at 16–17 pcw (n = 3 human samples/age). **(C)** SP neuron (green, eGFP transduced cell) with STXBP1 (red) present on neurites along thin putative axons extending from the cell body (i) and spine-like structures (ii) indicated by the white arrows. **(D)** Photomicrographs of a 16 pcw culture infected with AAV9-CMV-GFP and taken at 7, 21 and 27 DIV, showing the same neuron (highlighted by the masked images). Note the dynamic nature and elaboration of neurites over time. Coloured outlines highlight reference cells that are stable overtime.

### Synapse maturation in human cortical slice cultures

Given the relatively higher expression of STXBP1 in the subplate of 16–17 pcw tissue, we focused on this age for subsequent analysis of how the organotypic cultures continue to develop *in vitro.* To examine how neuronal morphology developed in the human cortical slice cultures, we infected cultures with viral vectors carrying the eGFP construct. We were able to resolve the neurites of individual labelled neurons and follow their development over intervals of many days to weeks, *in vitro*. Over these time periods, there was a marked elaboration of the neurite processes (Figure 1D).

The number of STXBP1 immunoreactive puncta in the subplate more than doubled, between 0 DIV and 27 DIV (16 pcw cultures increase from 4.1 ± 1.6 to 9.0 ± 0.4 puncta / 100 μm^3^; 17 pcw cultures increase from 4.2 ± 0.6 to 13.3 ± 1.1 puncta / 100 μm^3^; Figure 2A). Consistent with this, we also found significant increases in both glutamatergic synapses (vGlut1/Homer immunoreactivity; 16 pcw cultures increase from 2.7 ± 0.5 to 21.1 ± 1.7 puncta / 100 μm^3^; 17pcw cultures increase from 9.2 ± 0.9 to 30.3 ± 2.1 puncta / 100 μm^3^; Figure 2B), and GABAergic synapses (vGAT/gephyrin immunoreactivity; 16 pcw, increase from 1.0 ± 0.4 to 3.7 ± 0.6 puncta / 100 μm^3^; 17 pcw, increase from 0.6 ± 0.2 to 4.8 ± 0.4 puncta / 100 μm^3^; Figure 2C). These data show that in the human organotypic cultures, subplate neurons continue to mature, making both glutamatergic and GABAergic synapses.

**Figure 2.**
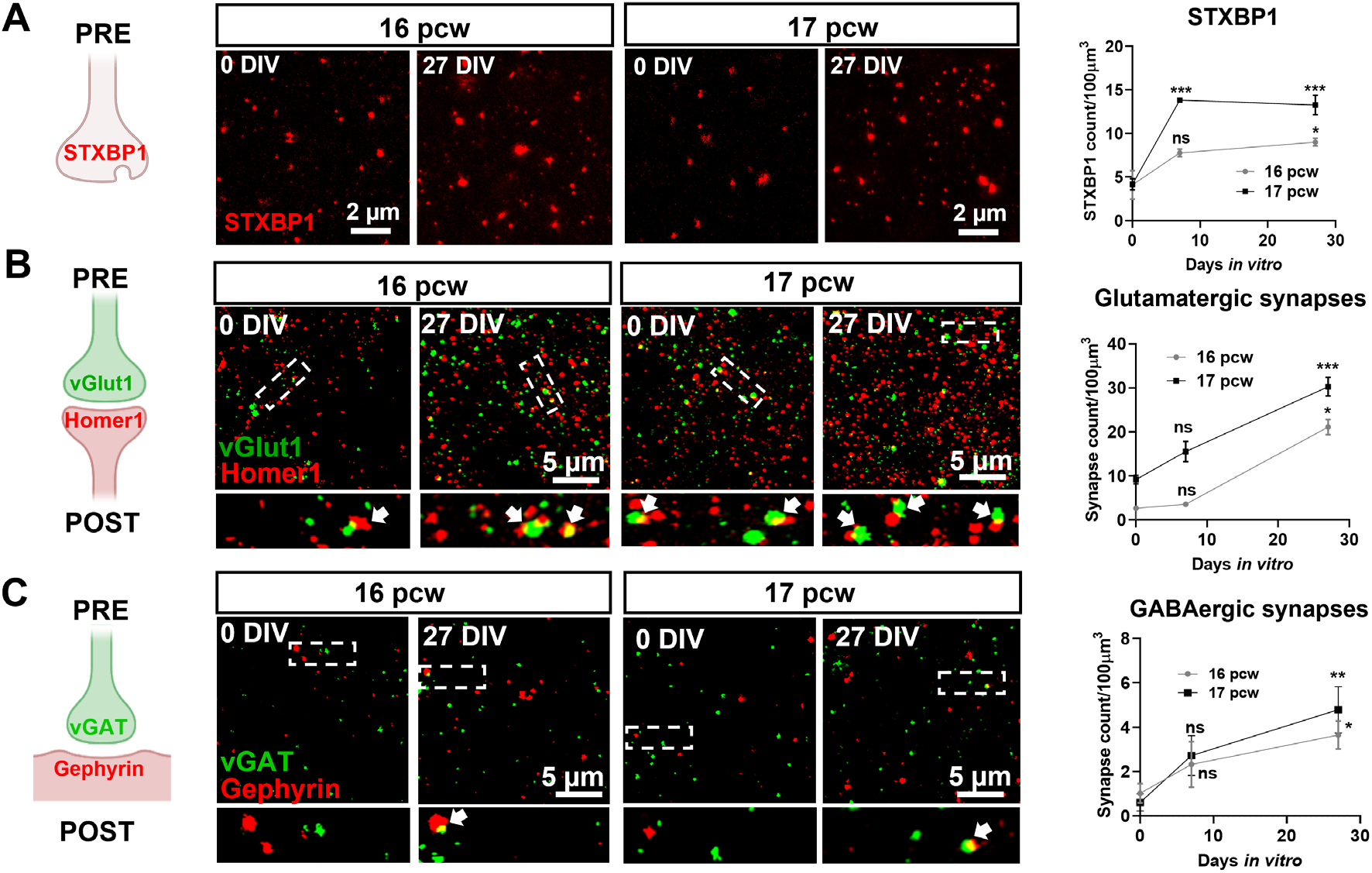
Synapse maturation in human cortical slice cultures. **(A)** Left: schematic displaying the pre-synaptic location of STXBP1. Middle: photomicrographs of STXBP1 (red) in the subplate of 16–17 pcw cultures at 0 and 27 DIV. Right: quantification of the increase in the number of STXBP1 puncta over time at both ages. **(B)** Left-middle: co-localisation of the pre- and post-synaptic glutamatergic markers, vGlut1 (green) and Homer1 (red), respectively indicated by arrows, in the subplate at 0 and 27 DIV. Areas, demarcated by dotted boxes in the upper micrographs, are shown enlarged below. All cultures were prepared at 16–17 pcw. **(C)** Similar analyses performed for GABAergic synapses based upon co-localisation of vGAT (pre-synaptic in green) and Gephyrin (post-synaptic in red). Analyses for panels (A) to (C) were performed as two-way ANOVA with repeated measures from comparisons made at 0 DIV for each age, n = 6–8 slices per time point from 2–3 human samples/age. Error bars indicate SEM.

### Divergent regulation of synapses with reduced STXBP1 levels

We next investigated the effects of manipulating the expression of STXBP1 in the human cortical slice cultures. We created a generic haplo-insufficiency phenotype for STXBP1 (“knock-down”) in our human cortical slice cultures using an AAV packaged *STXBP1* shRNA in 16 pcw tissue samples, derived from two individuals (Figure 3). To control for any effects of the viral infection, we performed similar infections with a scrambled STXBP1 sequence, in age matched brain slices (“scrambled control”). The shRNA achieved approximately 50% downregulation of STXBP1 at 14 DIV, compared with the scrambled control cultures prepared from the same brain tissue sample (Figure 3A). Within the subplate, STXBP1 immunoreactivity was reduced by approximately 30%, although there was no overall change in the absolute number of STXBP1 puncta (Supplementary Figure 1). Since each punctum was presumed to represent a synaptic terminal, this suggests that our shRNA design reduces endogenous STXBP1 protein levels, without affecting the total number of STXBP1-positive synaptic terminals.

**Figure 3.**
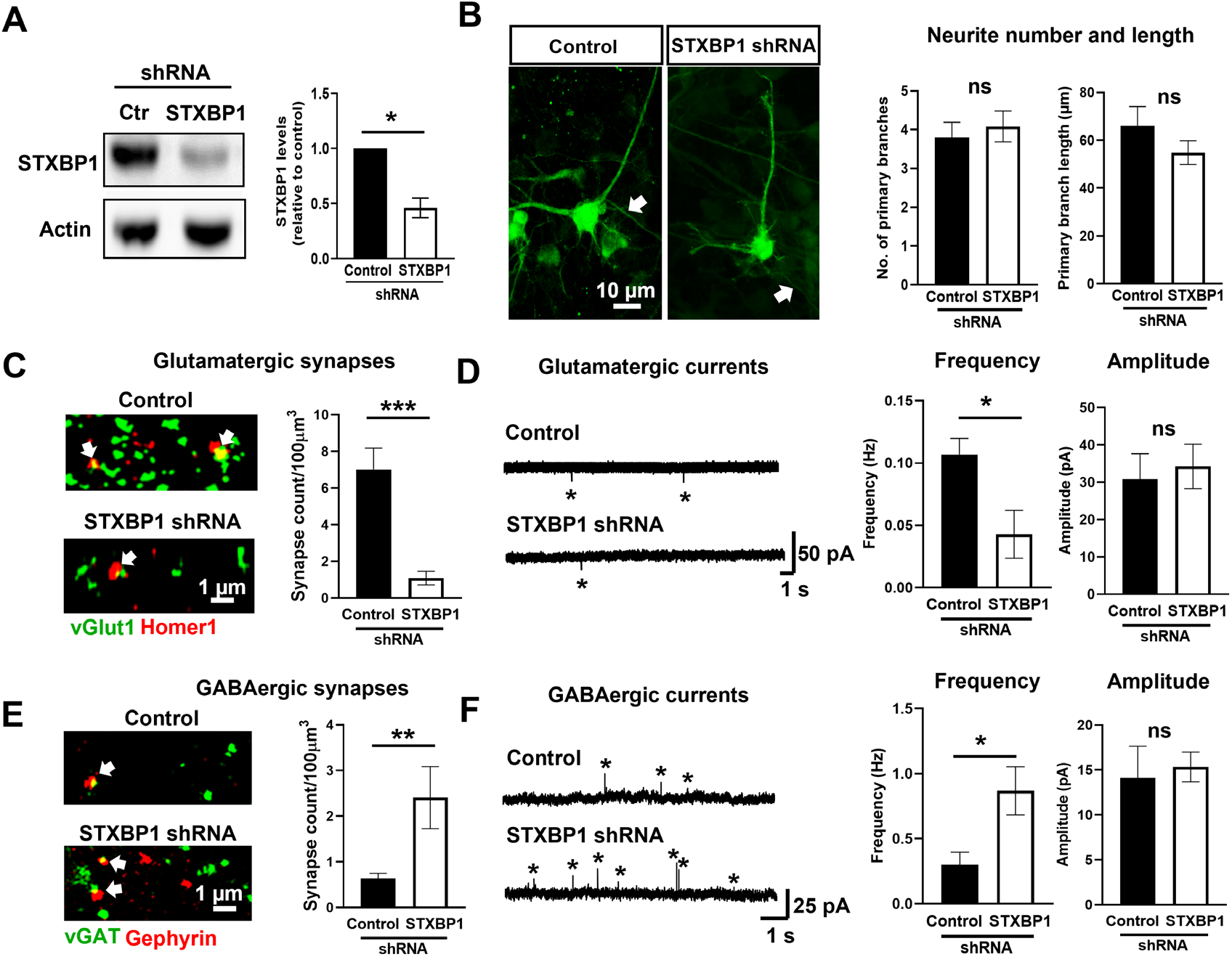
Divergent regulation of glutamatergic and GABAergic synapses in the subplate following knock-down of STXBP1. **(A)** Left: western blot showing knock-down of STXBP1 protein in a whole slice at 16 pcw following 2 weeks with scrambled control (Ctr) or *STXBP1* shRNA. Right: quantification of band intensity showing approximately 50% reduction in STXBP1 levels (unpaired t-test, taken from n = 5 slices/condition from 2 human samples). **(B)** Left: example subplate neurons infected with scrambled control or *STXBP1* shRNA for 2 weeks at 16 pcw. Note the putative axons highlighted with the arrows. Right: quantification of primary neurite number and length reveals no gross morphology changes in both conditions (n = 15 cells from 10 slices and 2 human samples). **(C-E)** Quantification of glutamatergic synapses (white arrows) (C) and GABAergic (E) synapses (white arrows) in the subplate at 16 pcw following two weeks of scrambled control or *STXBP1* shRNA by colocalisation of pre- and post-synaptic markers. Analysis performed using an unpaired t-test, n = 6–8 slices per time point from 2 human samples. (**D-F**) Representative miniature glutamatergic post-synaptic currents recorded at −60 mV (D) and miniature GABAergic post-synaptic currents recorded at 0 mV (F) from 16 pcw subplate cells in both genetic conditions. The frequency and amplitude of events (starred) were quantified (unpaired t-test, n = 8 cells per time point from 2 human samples). All data represented as mean ± SEM

We found no difference in the size of the subplate between control (n = 10 slices, average depth = 529 ± 25 μm) and knock-down cultures (n = 10, average depth = 549 ± 21 μm; p > 0.05, unpaired-test). Furthermore, the number and length of neurite processes in subplate neurons in STXBP1 knock-down cultures was indistinguishable after 14 DIV (Figure 3B). In contrast, in these same knock-down cultures, the number of glutamatergic synapses was reduced by approximately 86% within the subplate compared to cultures receiving scrambled control (Figure 3C). To evaluate the functional effects of these anatomical changes, we recorded glutamatergic currents. The frequency of synaptic events was decreased by 50% in cells expressing *STXBP1* shRNA (Figure 3D). Remarkably, the genetic manipulation had the opposite effect on GABAergic synapses in cultures undergoing STXBP1 knock-down, with nearly an 80% increase in number of GABAergic synapses within the subplate (Figure 3E), and a 70% increase in the frequency of GABAergic post-synaptic currents (Figure 3F), compared to scrambled control. There was no change, however, in the amplitude of either glutamatergic or GABAergic currents (Figures 3D and 3F). Overall, it appears that reducing STXBP1 expression has no effect on the overall stability of primary subplate neuron neurites but does alter synaptic stability during early subplate development.

### Impaired vesicle recycling with loss of STXBP1

STXBP1 plays a critical role in synaptic vesicle recycling^17, 18^. To assess this function, we assayed depletion of fluorescence in tissue that had been loaded with the lipophilic dye, FM4-64. Synaptic vesicles loaded with FM4-64 were identified along isolated axons in the subplate in 16 pcw tissue (Figure 4A). Under baseline conditions, signal intensity was reduced by 30% from 0 to 12 seconds in scrambled controls but not after *STXBP1* shRNA knock-down (Figure 4B), indicating an impairment in spontaneous neurotransmission at synapses with reduced STXBP1 levels.

**Figure 4.**
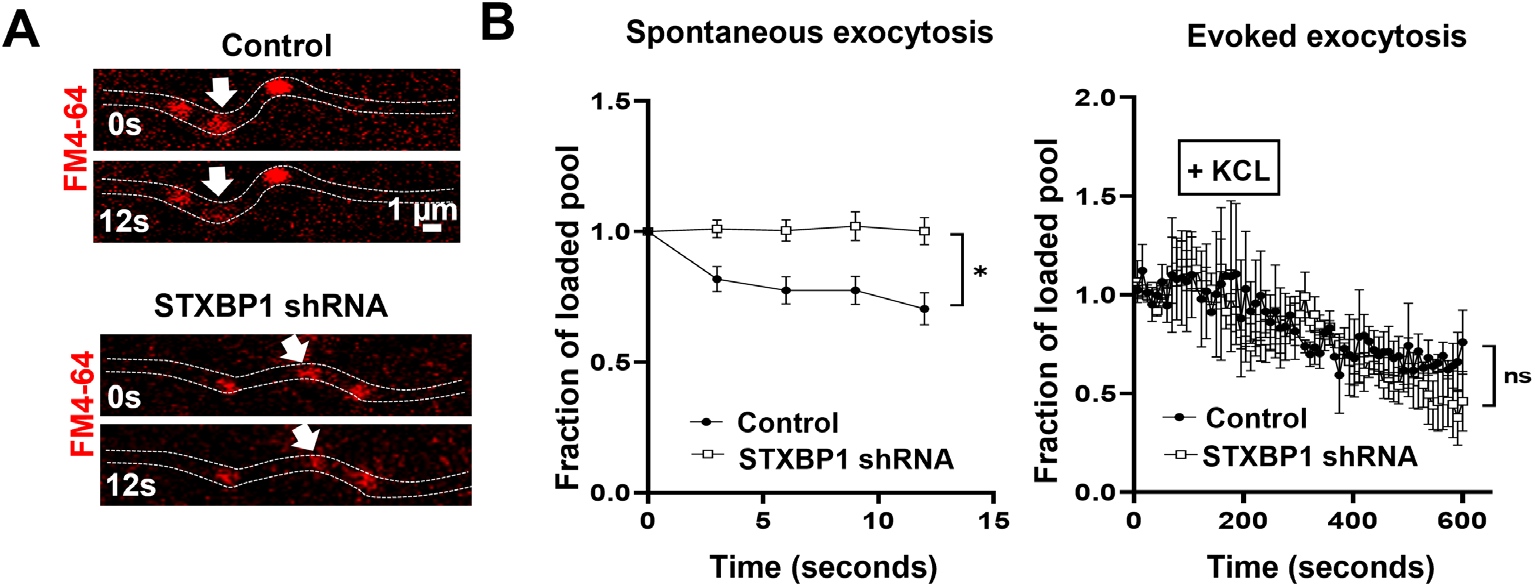
Impaired spontaneous vesicle release with reduced STXBP1 levels. **(A)** Isolated axons tracked from FM4-64 loaded subplate neurons in scrambled control and STXBP1 shRNA slices (16 pcw). Note the presence of spontaneous release from vesicles loaded with FM4-64 (white arrows). **(B)** Quantification of spontaneous exocytosis (left; n = 8 slices from 2 human samples) and evoked exocytosis (right; n = 6 slices from 2 human samples) following brief potassium chloride (KCl) stimulation in both genetic conditions. Analysis was performed using a two-way ANOVA with repeated measures. All error bars represented as SEM.

To assess evoked exocytosis, slices were additionally perfused with a depolarisation inducing concentration of potassium ions. The intensity of the signal started to decrease 120 seconds after the start of stimulation until the end of the experiment (overall 50% loss) in both scrambled controls and *STXBP1* shRNA, but there was no difference between the experimental groups (Figure 4B). This indicates that reduced STXBP1 does not affect evoked exocytosis in the subplate under these experimental conditions.

## Discussion

We present a novel means of investigating the effect of genetic disorders on prenatal human cortical development that benefits from retaining subplate and cortical plate networks. This represents a major advance over other approaches, such as human cerebral organoid or induced pluripotent stem cell cultures, neither of which show structures that approach the complexity of the real *in utero* networks. In these initial proof-of-principle studies, using a simple shRNA interference genetic manipulation, we demonstrate the divergent effects of STXBP1 haploinsufficiency on glutamatergic and GABAergic neurons at an early stage of brain development. This work will help improve our understanding of the pathological mechanisms underlying certain early infantile onset developmental and epileptic encephalopathies^11^.

Reduced levels of STXBP1 do not affect the gross morphology of subplate neurons at early developmental stages. Previous studies have shown that absence of *UNC-18,* the ortholog gene of *STXBP1,* does not affect neurite length or maintenance in C. elegans ^18^ and that heterozygous Stxbp1 haploinsufficiency does not affect dendrite length^17^ or cortical neuron density^11^ in mice postnatally. By addressing these questions in a model closer to the human brain, our data provides confirmatory evidence that these findings are relevant also to the human disorder.

Deficiencies in STXBP1 can result in glutamatergic synaptic neurotransmission dysfunction in rodent models^11^ and neurons derived from human embryonic stem cells^19^. Similarly, we find reduced glutamatergic post-synaptic current frequency and synapse number in the human subplate following knock-down of STXBP1. Interestingly, we observed the opposite effect on GABAergic synapses, with an increase both in the frequency of GABAergic synaptic events, and the density of GABAergic synapses. STXBP1 is normally present at both GABAergic and glutamatergic synapses, so the divergent effects of STXBP1 knock-down on the two types of synapses was unexpected; given that the two synaptic classes are likely to be regulated in parallel, this divergence may represent a compensatory change manifest in one class, responding to a primary effect in the other. This new finding demonstrates an advantage of our model over previous single cell assays that may not be sensitive to such divergent effects present in an intact neuronal network.

The large decrease in frequency of glutamatergic currents, without a change in amplitude, is indicative of an impairment in pre-synaptic neurotransmitter release. Reliable and sustainable neurotransmitter release is essential for effective neuronal communication. Both spontaneous, and evoked neurotransmitter release is affected by lowering the levels of STXBP1 in many animal models^17–20^. These results are consistent with the observation that at mature synapses STXBP1 is crucial for the assembly of the SNARE complex, a collection of proteins involved in the fusion of vesicles with the plasma membrane^21–23^, variants in several of which are implicated in neurodevelopmental disorders and epilepsy (collectively, termed SNAREopathies^9^). Conversely, we observe impairments in spontaneous neurotransmitter release, but not evoked exocytosis. Interestingly, a similar dissociation between impaired spontaneous, but normal evoked release has previously been reported in mouse neurons expressing the human disease variant C180Y^11^. Our results may reflect some compensation from other cooperative proteins known to function together to chaperone SNARE assembly, including MUNC13-1^24, 25^. STXBP1 could also have a more important role in spontaneous than evoked exocytosis at early developmental stages, because spontaneous neuronal activity is the predominant driving force controlling network formation^26^.

In summary, we present a new experimental approach for understanding human cortical development at a stage approximately equivalent to 14-18 pcw. Loss of STXBP1 at this stage has divergent effects upon glutamatergic and GABAergic synapses in the subplate, reducing the former, but increasing the latter. The synaptic networks in this region are important for neurogenesis, neuron migration, and formation of thalamocortical and cortico-cortical connectivity^4, 27, 28^. Therefore, alterations in subplate synaptic activity are expected to have a large effect on the subsequent development of the overlying cortex. Treatment of STXBP1-related disorders is challenging, requiring a personalised treatment plan^10^. This model system could serve as a crucial bridge to facilitate the development of future precision-medicine approaches.

## Abbreviations

HDBR: Human Developmental Biology Resource
STXBP1: Syntaxin Binding protein 1
MUNC18-1: Mammalian uncoordinated-18
pcw: post conception week
DIV: days *in vitro*
AAV: adeno-associated virus
shRNA: short hairpin RNA

## Acknowledgements

This work would not have been possible without the tissue donation to the HDBR and the efforts of everyone within the biobank and the help of the Bioimaging Unit at Newcastle University.

## Author contributions

F.M. performed all experimental procedures in the manuscript, design of experiments and conceptualising the project. A.D. assisted with the immunohistochemistry and image analysis. D.L-S. and R.H.T. were involved with conceptualisation of the project. F.M., A.J.T. and G.J.C. conceived the overall project, helped design the experiments and analyses. Funding for the project was obtained by A.J.T. All authors participated in the interpretation of data and writing of the manuscript.

## Supplementary Figures

**Supplementary Figure 1:**
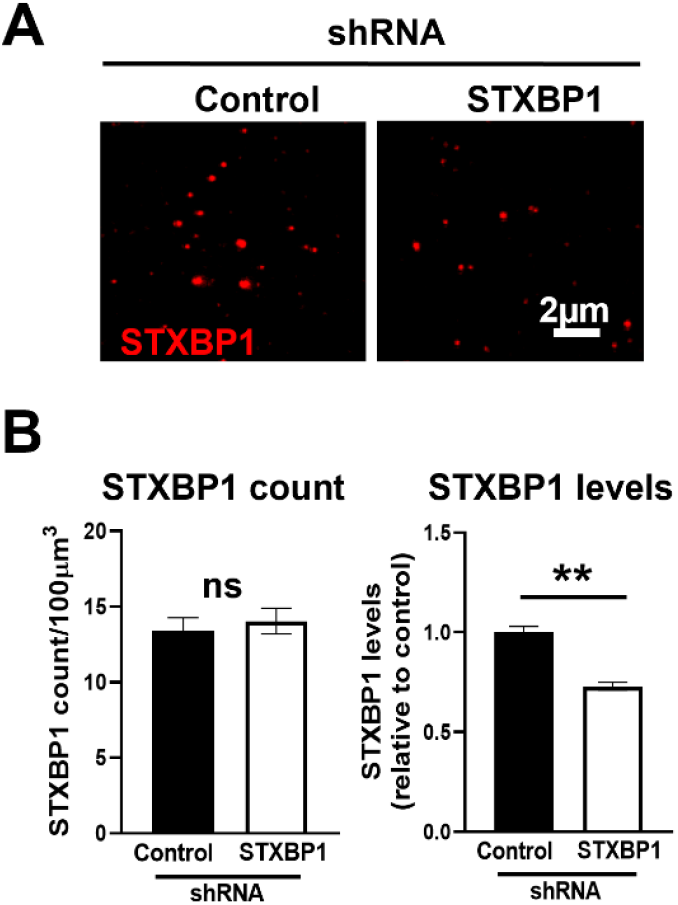
Genetic knock-down of STXBP1 in the subplate of human cortical slice cultures. **(A)** Photomicrographs of STXBP1 (red) in the subplate in the subplate at 16pcw following 2 weeks with scrambled control or *STXBP1* shRNA **(B)** Quantification of STXBP1 puncta count and integrated density levels in the subplate. Note the comparable terminal numbers (STXBP1 count) but less protein (STXBP1 levels). Analysis performed using an unpaired t-test, *n = 8* slices/condition from *2* human samples). All data represented as mean ± SEM.

## Supplementary Methods

### Human organotypic slice cultures

Tissue was transferred to oxygenated ice-cold high sucrose ACSF containing (in mM): 180 sucrose, 25 NaHCO_3_, 10 glucose, 10 MgSO_4_, 2.5 KCl, 2 N-acetylcysteine, 1.25 NaH_2_PO_4_, 1 taurine, 1 ascorbic acid, 0.5 CaCl_2_, 0.1 aminoguanidine hydrochloride, 0.044 indomethacin, 0.044 ethyl pyruvate and adjusted to 300–310 mOsM and 7.4 pH. Subsequently 280μm slices were cut using a 5100mz Vibratome (Camden Instruments). Cortical slices were placed inside 6-transwell plates containing BrainPhys (5790; Stemcell Technologies) culture media supplemented with 1 × N2, 1 × B27, 40ng/ml Brain-derived Neutrophic Factor (BDNF), 20ng/ml Glia-derived Neurotrophic Factor (GDNF), 30ng/ml Wnt7a, 200 nM ascorbic acid, 1 mM dibutyryl cyclic AMP and 1 μg/ml laminin.

### Live imaging

Live imaging was performed using an inverted Zeiss LSM800 Airyscan confocal microscope. AAV9-CMV-GFP infected slices were placed in culture media and maintained at 37°C, 5% CO_2_. Images were acquired in the subplate using a x20 objective (Numerical Aperture; NA = 0.8) every 30 seconds for 15 minutes from the same slice for 4 weeks generating final image resolutions of 512 × 512 pixels.

FM4-64 dye was utilised as previously described^14^. Briefly, infected slices were transferred to control solution containing (in mM, pH 7.3): 120 NaCl, 10 HEPES, 10 Glucose, 2 CaCl_2_ and 1 MgCl_2_ to equilibrate. After 5 minutes, they were placed in control solution, FM4-64 dye (7.5 μM) and potassium chloride (KCl; 50 mM with osmolarity compensated) for 90 seconds to stimulate and load the dye in vesicles. Subsequently, slices were placed for 60 seconds in control solution and FM4-64 dye. Finally, slices were transferred to control solution to wash off any excess non-internalised dye.

Consecutive images were acquired of FM4-64 loaded vesicles using a 40x objective (NA = 1.3) at room temperature (RT). Slices were firstly perfused with control solution and axonal regions loaded with FM4-64 vesicles identified for imaging. After 10 minutes, vesicle exocytosis (FM4-64 dye unloading) was stimulated by perfusion with control solution (substituted with 50mM KCl) for 150 seconds, before reperfusion with control solution for a further 3 minutes. Simultaneous images of the GFP-positive axons and FM4-64 loaded vesicles were acquired just before, during and after stimulation, with a 3 second delay between each frame.

Analysis was performed using Fiji software (NIH, RRID:SCR_002285). Regions of interest (ROIs) representing the vesicles, were identified for each slice. One ROI on an area with no signal (representing the background) and one ROI on an area with a stable signal (to account for the FM4-64 bleaching) were selected. The ROIs remained the same throughout the videos and their mean fluorescent measurements were quantified. Mean background values were subtracted from the measured signal, and each value was divided by the mean stable signal value. Finally, these values were normalised to the average ROI values of all frames 60 seconds before stimulation to calculate the fraction of loaded vesicle pool. Evoked exocytosis was averaged every 9 seconds. For spontaneous exocytosis (just before stimulation), ROI values were normalised to the average of first five frames.

### Whole cell-patch clamp recordings

Slices were transferred into the recording chamber of an upright Leica DMLFSA fluorescent microscope fitted with Micro Control Instruments micromanipulators and continuously perfused at 34°C with oxygenated ACSF containing (in mM): 126 NaCl, 3 KCl, 1.25 NaH_2_PO_4_, 24 NaHCO_3_, 10 Glucose, 1.2 CaCl_2_ and 1 MgSO_4_. Cells were voltage-clamped in whole cell configuration using borosilicate glass patch electrodes (5–8 MΩ) filled with an intracellular solution containing (in mM): 125 K-methyl-SO_4_, 10 HEPES, 2.5 Mg-ATP, 6 NaCl, 290 mOsM and pH 7.35. All data were collected using an Axopatch 200B amplifier, filtered (1 kHz) and digitised (10 kHz). Miniature currents were monitored and analysed using WinEDR and WinWCP software (http://spider.science.strath.ac.uk/sipbs/software_ses.htm).

### Immunohistochemistry

Slices were fixed in 4% paraformaldehyde (PFA)/4% sucrose in phosphate buffered saline (PBS; 0.12 M, pH 7.4; PBS) for 20 minutes at room temperature (RT). Staining was performed using the protocol optimised specifically for acute slices^15^. In brief, slices were incubated in blocking solution (10% donkey serum, 1% Triton X-100 in PBS) for ~ 4-6 hours at RT on a vibrating shaker. Primary antibodies against Synapsin1 (1:500; Millipore, AB1543P), vGlut1 (1:2000; Millipore, AB5905), Homer1 (1:1000; Synaptic Systems, 160006)), vGAT (1:1000; Synaptic Systems, 131003), Gephyrin (1:500; Synaptic Systems, 147318), NeuN (1:1000; Abcam, AB104224), STXBP1 (1:500; Abcam, ab109023), GAD67 (1:1000; Millipore, MAB5406), MAP2 (1:2000, Abcam, AB5392) or GFP (1:5000; Abcam, ab290) were applied overnight at 4°C. Slices were subsequently incubated with fluorophore-bound secondary antibodies (1:500; Life Technologies) for 2-3 hours at RT on a vibrating shaker before DAPI (1:5000) applied and mounted on glass slides in Fluoromount-G (SouthernBiotech).

### Image analysis

The number as well as the length of primary subplate neuron processes was measured using Fiji software. Synaptic counts and integrated density (area × mean intensity) measurements were conducted using Zen 3.1 software (Zeiss; RRID:SCR_013672). These were optimised with a range of filters and thresholds to specifically identify the ROI for pre- and post-synaptic markers. Settings for markers of interest were kept consistent throughout all replicates. A synapse was defined as the colocalisation of specific pre- and post-synaptic markers with a minimum of 2 pixels overlapping. The counts of individual and colocalised ROIs were normalised to the volume of each image then represented per 100 μm^3^.

### Western blot

Membranes were blocked then probed overnight at 4°C on a vibrating shaker with anti STXBP1 antibody (1:1000; Abcam) and anti β-actin (1:5000; Cell Signalling). Appropriate species specific HRP-conjugated secondary antibodies were added (BIO-RAD) at RT. Visualisation of protein bands was performed using a GBOX-Chemi-XRQ gel system (Syngene).

